# A new immunocompetent rectal cancer model to study radiation therapy

**DOI:** 10.1101/2022.02.07.479335

**Authors:** Jin K. Kim, Chao Wu, Michael Del Latto, Yajing Gao, Seo-Hyun Choi, Maria Kierstead, Charles-Etienne Gabriel Sauvé, Canan Firat, Almudena Chaves Perez, Jussi Sillanpaa, Chin-Tung Chen, Kayla E. Lawrence, Philip B. Paty, Francisco M Barriga, Jinru Shia, Charles L. Sawyers, Scott W. Lowe, Julio García-Aguilar, Paul B. Romesser, J. Joshua Smith

**Author notes:** Co-corresponding authors |.

## Abstract

We describe a new mouse model of rectal cancer (RC) involving rapid tumor organoid engraftment via orthotopic transplantation; the resulting RC tumors invaded inward from the mucosal surface and metastasized to distant organs. Histologically the tumors closely resemble human RC and mirror remodeling of the tumor microenvirnoment (TME) in response to radiation. This murine RC model thus recapitulates the pathogenesis of human RC, thereby fulfilling the need for a physiologically accurate model for preclinical efficacy studies.

## Main

Rectal cancer (RC) constitutes approximately one-third of colorectal cancer diagnoses and increasingly affects younger individuals (<50 years old)^1^. RC, unlike colon cancer, is typically treated with radiation therapy (RT) and sometimes chemotherapy before radical surgery. Growing evidence suggests that the immune tumor microenvironment (TME) plays crucial roles in tumorigenesis and response to treatment^2–4^. In turn, RT, by inducing DNA damage and cell death, may induce TME remodeling^5^ and influence immunological responses^4,6,7^. Investigating these ideas will require immunocompetent murine models that accurately mimic human RC and its response to RT.

Numerous murine models of colon cancer have been established, but these do not offer the ideal setting in which to examine the TME for RC as the pelvic anatomy and blood supply are unique. A few models have been developed in the proper rectal anatomic context, but these involve either chemical induction, which can cause chronic colitis and alter the TME, or intrarectal injection of tumor cells, which results in submucosal-initiated tumors, unlike human tumors which initiate in the mucosa^8,9^. The advantages and limitations of existing models are summarized in **Supplementary Table 1**.

An additional need is optimized RT delivery methods for murine RC models. Many studies have used whole-body radiation, but this is not representative of localized pelvic RT in patients and adds undesirable systemic effects.^10,11^. To address these needs, we sought to develop an anatomically and physiologically accurate RC model, along with a pelvic irradiation method.

We isolated and cultured organoids from genetically engineered (*Apc*^*flox/flox*^, *LSL-Kras*^*G12D/+*^, *Tp53*^*flox/flox*^) mouse (C57Bl/6J) rectal tissue. Organoids were transduced ex vivo with Adeno viral vector expressing Cre-recombinase to activate Kras and inactivate Apc and p53 (*Apc*^*-/-*^, *Kras*^*G12D/+*^, *Tp53*^*-/-*^; hereafter referred to as AKP). This represents a common genetic configuration in human RC. Organoids of the desired genotype were selected by removing Wnt ligands and EGF from the media, followed by Nutlin-3 selection^8^.

The resulting AKP rectal tumor organoids (tumoroids) were prepared as rectal enema. Mice (C57Bl/6J) were anesthesized and a small-caliber brush was inserted through the anus to mechanically disrupt the rectal mucosa (**Fig. 1a-b**). Thereafter, tumoroids were transanally instilled and the anus was glued shut for 6-8 hours to prevent leaakage and facilate tumoroid engraftment. Step-by-step instructions are in **Supplementary Video 1**. Of note, this procedure is faster than other orthotopic tumor implantation procedures and requires minimal to no preparation in advance. It is important to note that mechanical disruption reduces anesthesia time and animal stress compared to DSS methods allowing more animals to be engrafted in a less invasive manner. Tumoroids engraft at the site of mucosal disruption (**Fig. 1c**) and preserve WNT and KRAS pathways alterations *in vivo*, as demonstrated by elevated β-catenin and phospho-ERK (**Supplementary Fig. 1**).

**Figure 1.**
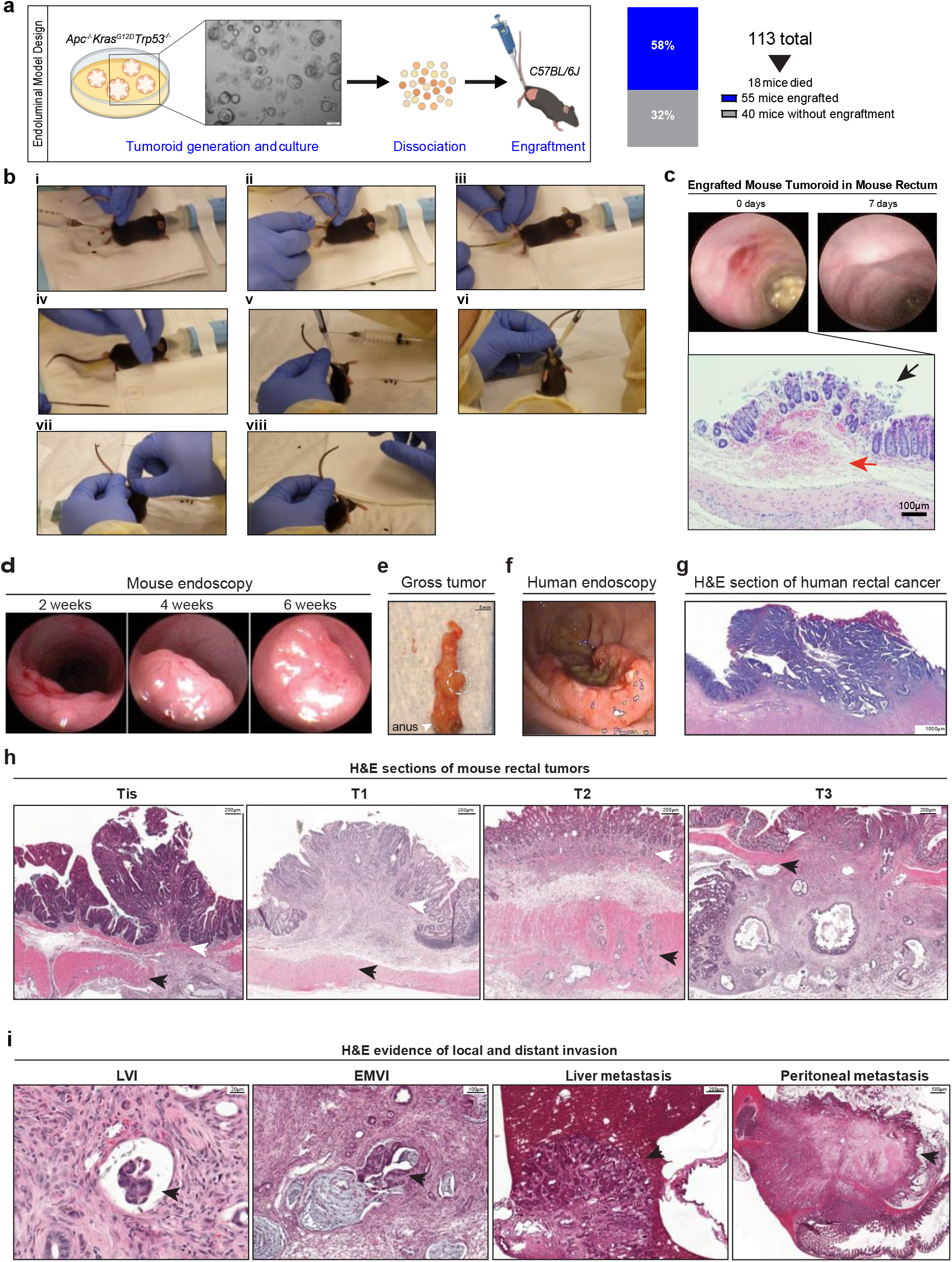
A new orthotopic rectal cancer model in immunocompetent mice. **a**, Left panel: diagram of rectal tumoroid preparation. AKP rectal organoids were isolated from a transgenic mouse, cultured in Matrigel, and then harvested and transferred as an enema to C57BL/6J mice after mechanical disruption of the rectal mucosa. Right panel: rates of post-procedure mortality and engraftment. **b**, Critical steps in endoluminal engraftment. i) irrigate rectal vault with PBS. ii) insert a trimmed P200 pipette tip into the anus as a guide. iii) insert and irritate mucosa with a small-caliber brush. iv) confirm a blood smear on the brush, which indicates adequate endoluminal disruption. v) pipette tumoroid mixture into the rectal vault. vi) seal the anus with Vetbond glue. vii) pinch anus closed. viii) allow Vetbond to dry and confirm anus is sealed. **c**, Tumoroids implanted at the site of mucosal disruption. Endoscopy immediately after brush and 1 week after engrafting shows tumoroids implanting at the site of mucosal disruption. **d**, Serial endoscopy pictures of the engrafted tumoroids post-transplantation. **e**, Necropsy shows tumor (circled) in the distal rectum. **f**, A representative endoscopic image of a human rectal cancer. **g**, H&E section of human rectal cancer showing adenocarcinoma invading from the mucosa. **h**, H&E sections of engrafted tumors in this murine model. Tumors ranging from carcinoma in situ (Tis) to T3 are presented. White arrows point to the muscularis mucosa and black arrows to muscularis propria. **i**, H&E sections showing local invasion (lymphovascular invasion [LVI] and extramural vascular invasion [EMVI]) and distant metastasis to liver and peritoneum of engrafted tumors. Black arrows point to the tumor.

Of 113 mice treated, 18 (16%) died within 2 days after transplantation due to intestinal perforation (confirmed at autopsy). Post-procedural mortality within 36 hours decreased as experience with the technique increased. Among the 95 surviving mice, this procedure yielded a 58% engraftment rate (**Fig. 1a, right panel**). Tumor engraftment (seen by either a luminal mass or increasing size of a luminal lesion) was apparent on colonoscopy by 2-4 weeks after transplantation (**Fig. 1d**). The median distance of the tumor from the anal verge was 3.5 mm (range, 0-10 mm) (**Fig. 1e**). Engrafted tumors grew and obstructed the rectal lumen within 1-3 months of transplantation (**Fig. 1d**), and were macroscopically similar to human RC under endoscopy (**Fig. 1f**). Moreover, histopathologic review showed that the engrafted tumors resemble the development and progression from pre-invasive to invasive cancer and to metastatic disease (**Fig. 1h-i**) as observed in human RC (**Fig. 1g**). These data establish the feasibility of this RC model and its clinically relevant patterns of tumor progression and metastasis.

To test the application of our model to RT, we applied localized pelvic RT to tumor-bearing mice using a customized cerrobend apparatus (**Fig. 2a-b**). The experimental design is illustrated in **Fig. 2c**. The mice were irradiated when the engrafted tumors had grown to encompass 20-30% of the rectal lumen (**Supplementary Fig. 2**), which occurred at a median of 4.5 weeks after tumor cell transplantation. The mice were then randomized to RT (15-Gy single dose; RT group) or no treatment (control group). All mice were monitored carefully for health status and subjected to bi-weekly surveillance endoscopy. Mice were euthanized when the circumferential tumor involvement exceeded 50% of the rectal lumen.

**Figure 2.**
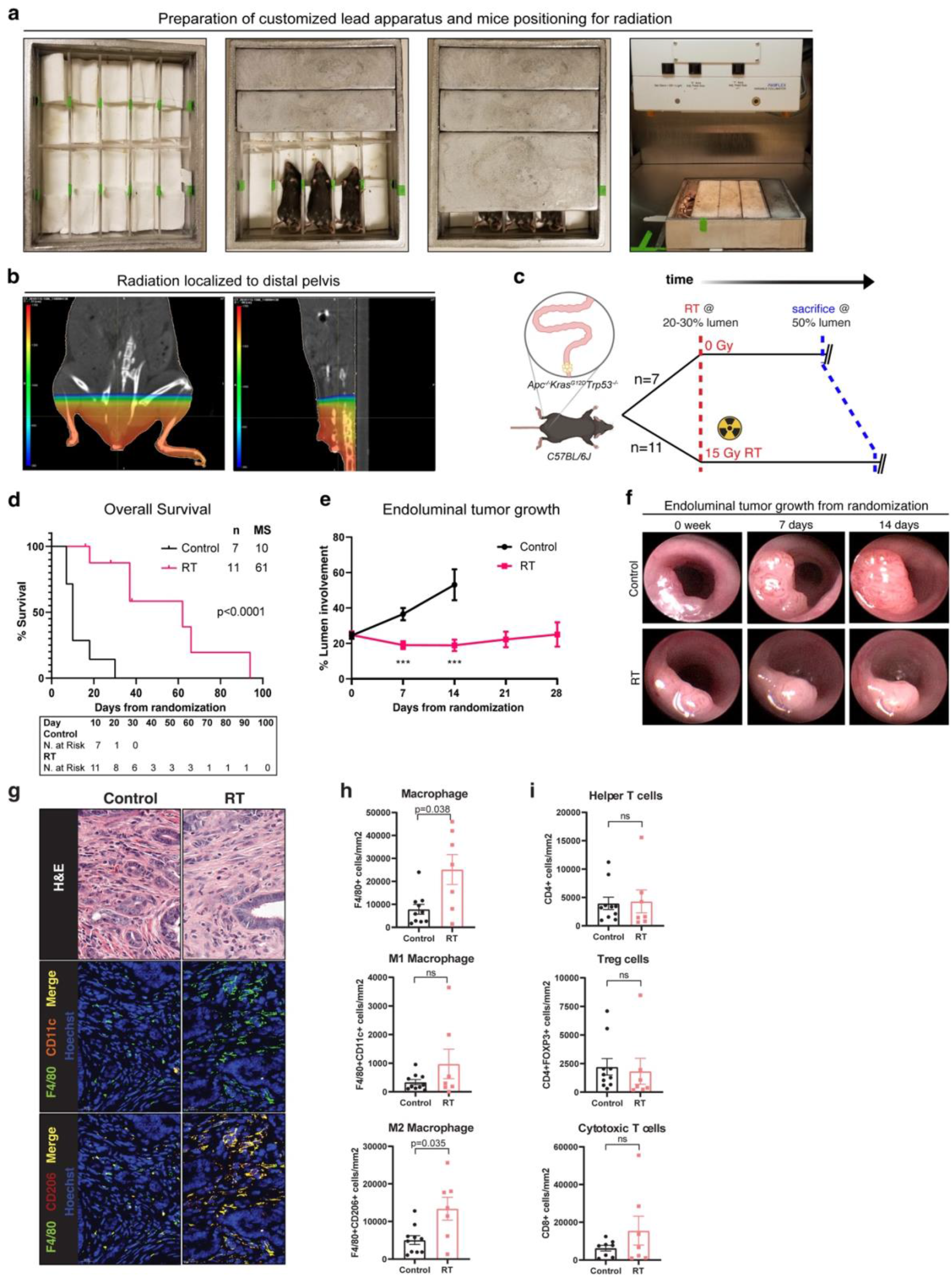
Application and evaluation of localized radiation therapy in this mouse model. **a**, Set-up to provide localized rectal radiation to mice using customized cerrobend blocks. **b**, Radiation dosimetry of the predicted radiation field to the distal pelvis. **c**, Study schema, including randomization of mice to the control group (n=7) or 15 Gy of RT group (n=11). The beam-on time was calculated using the dose rate to air and does not incorporate corrections for absorption and scatter in the mouse. **d**, Kaplan-Meier analysis of overall survival of control and RT group mice. Mice that were sacrificed for having tumor involvement > 50% of the lumen were deemed as events. Log-rank p-value is shown. **e**, Mean circumferential tumor size of the control and RT group mice at the time of treatment and at weeks 1-4 from treatment are plotted. Error bars denote the standard error of mean. ***p<0.0001 by unpaired two-tailed t-test. **f**, Representative serial endoscopic pictures of a control mouse and an irradiated mouse. **g**, Top: representative hematoxylin & eosin (H&E) stained images of rectal tumors from control mouse and irradiated mice. Middle and bottom panels: representative images of immunofluorescent staining for tumor-associated macrophages. F4/80 (green) is a macrophage marker; CD11c (orange) was considered to be a marker for M1 polarized macrophages and CD206 (red) for M2 polarized. Counterstain with Hoechst (blue). **h**, Quantification of tumor-associated macrophages and the M1/M2 subtypes. Error bars denote the standard error of mean. P-values provided by unpaired two-tailed t-test. ns, p-value > 0.05. **i**, Quantification of CD4^+^ helper T cells, CD4^+^FOXP3^+^ Treg cells, and CD8^+^ cytotoxic T cells. Error bars and p-values are as in panel h.

Mice tolerated RT well without significant weight loss or illness. At one week post-RT, peripheral blood showed a decrease in all five types of white blood cells (although statistical significance was not reached due to small sample size)—indicating systemic leukopenia in RT-treated mice (**Supplementary Fig. 3**). This is consistent with the acute transient hematological toxicity seen in patients^12^. We also compared the survival rate between pelvic-irradiated mice and whole body irradiated mice to ensure our device provided effective bone marrow shielding and mitigated lethality from whole body irradiation (**Supplementary Fig. 4**).

RT resulted in a 10-40% treatment effect upon tumor regression grading by histolgical examination. This treatment effect is similar to what we see in RC patients^13^. Interestingly, tumors initially decreased in size, as assessed endoscopically, after RT, but most eventually regrew (**Fig. 2e**). RT resulted in significantly better survival (**Fig. 2d**) and reduced tumor growth (**Fig. 2e-f**).

We next examined changes in the TME to evalaute if our model recapatituates changes in the TME in response to RT in human patients. Pior work has show a significant increase in M2 macrophage polarization post RT^14^. Similarly, when examining the TME of mice with AKP endoluminal tumors we observed a signfiant increase in macrophage infiltration (detected by macrophage surface marker F4/80) and M2 polarization (CD206^+^ F4/80^+^), but not M1 (CD11c^+^ F4/80^+^) (**Fig. 2g-h**). These data demonstrate that our model recapitulates the human tumor immune microenvironment and its response to RT.

In summary, we present a new murine model that recapitulates the tumor biology of RC and shows clinical relevance in response to RT. This model has a wide range of potential preclinical applications, such as the evaluation of radiosensitizers, combinations of RT with immunomodulators, and novel therapies. Furthermore, this model can be used to explore how radiation resistance is affected by various oncogenic mutations. For any of these applications, our method provides rapid generation of localized, mucosa-initiated rectal tumors in a model that is reliable, reproducible, and translatable to human RC.

## Methods

Step-by-step protocols and details on reagents and equipment are provided in **Supplementary Information**.

### Mice

We used 6- to 8-week-old female C57Bl/6J mice (Jackson Laboratory, stock no. 000664). All animal experiments were conducted under protocols approved by our institution’s Institutional Animal Care and Use Committee (11-06-012 and 06-07-012).

### Culturing and preparation of tumoroids for endoluminal transplantation

The AKP tumoroids were derived and cultured as described^8^. Rectal tumoroids were prepared for endoluminal transplantation as described^15^. Briefly, for endoluminal transplantation, the tumoroids were released from Matrigel, then resuspended in ice-cold PBS with 5% Matrigel to a concentration of 2×10^5^ cells/50 µL per mouse.

### Mechanical disruption of the mouse rectal mucosa and tumoroid transplantation

Critical steps are shown in **Supplementary Video 1**. Briefly, anesthesized mice (2% isoflurane and oxygen) underwent rectal irrigation to liberate stool. A smooth-trimmed P200 pipette tip (lubricated with Vaseline) was inserted into the anus as guide, then a small-caliper brush was inserted through the pipette tip and moved gently in and out of the rectum 3-5 times to gently disrupt the rectal mucosa. The tumoroid suspension was then slowly injected transanally using a P200 pipette. The anus was sealed using 5-10 µL of Vetbond Tissue Adhesive to prevent luminal contents from spilling. This bond was removed 6-8 hours later.

### Tumor surveillance and measuring tumor growth

Tumor engraftment was monitored by 1.9-mm rigid 30° small animal endoscope at least weekly. A video of each endoscopy was taken, and a static picture of the lesion was analyzed to calculate endoluminal involvement. Tumor growth was quantified by the percent of the field of view occupied by the tumor area as described^15^.

### Localized pelvic radiation

Mice were anesthetized by intraperitoneal injection of Ketamine/Xylazine (100 mg/mL; 10 μl/g body weight). Localized pelvic irradiation was delivered using an X-Rad 320 machine (Precision X-Ray, Madison, CT) (250kVp/ 12mA) with customized cerrobend blocks (fabrication technique will be shared upon request).

### Histopathology

Dissected tumor samples were fixed with 4% paraformaldehyde, embedded in paraffin, and sectioned according to standard protocols. For histopathologic evaluation, 5-µm sections were stained with H&E.

### Immunofluorescence

Tissue sections were deparaffinized, then boiled in pH 6.1 citrate buffer for 20 minutes for antigen retrieval. Sections were blocked in 10% normal donkey serum and 1% bovine serum albumin at room temperature for 1 hour, immunostained overnight at 4°C with primary antibodies, then for 2 hours at room temperature with fluorophore-conjugated secondary antibodies. Antibodies are listed in **Supplementary Information**. Cell nuclei were labeled with Hoechst dye.

### Imaging

Slides were scanned using the Pannoramic Flash slide scanner using a 20 × 0.8 NA objective. Images were examined and representative areas exported using CaseViewer 2.2. No gamma changes were made to any immunofluorescence images. All brightfield images are unaltered. Immune cells in total tumor regions were quantified using custom macros written in Image J (NIH, Bethesda, MD, USA). The number of immune cells in 1mm^2^ of tumor regions were caculated from each sample and averaged per group, and a Student’s t-test was performed for statistical analysis.

### Complete blood count of peripheral blood

Before endpoint collection, blood was collected in BD Microtainer tubes with K_2_EDTA and mixed on a Stuart Roller Mixer SRT9D. Complete blood counts were obtained using a Hemavet 950.

## Figure Legends

**Supplementary Information** includes **Supplementary Tables, Supplementary Figures**, and **Supplementary Note**.

**Supplementary Video 1**. Video illustrating the steps of the endoluminal tumoroid engraftment procedure

