## Supplementary Information for "A new immunocompetent rectal cancer model to study radiation therapy"

### Supplementary Figures

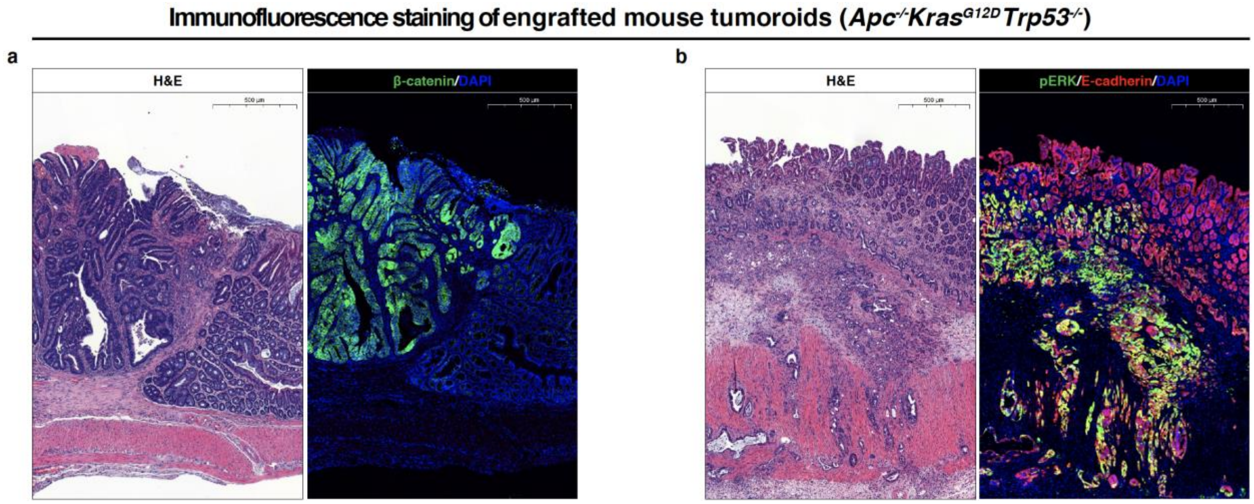


**Supplementary Figure 1. Serial sections of H&E and immunofluorescence staining of engrafted AKP rectal tumoroids**. (**a**) left: H&E, and right: β-catenin staining (green), counterstain with DAPI (blue). (**b**) left: H&E, and right: phospho-ERK staining (green), counterstain with DAPI (blue); E-cadherin (red) was used as an epithelial cell marker.

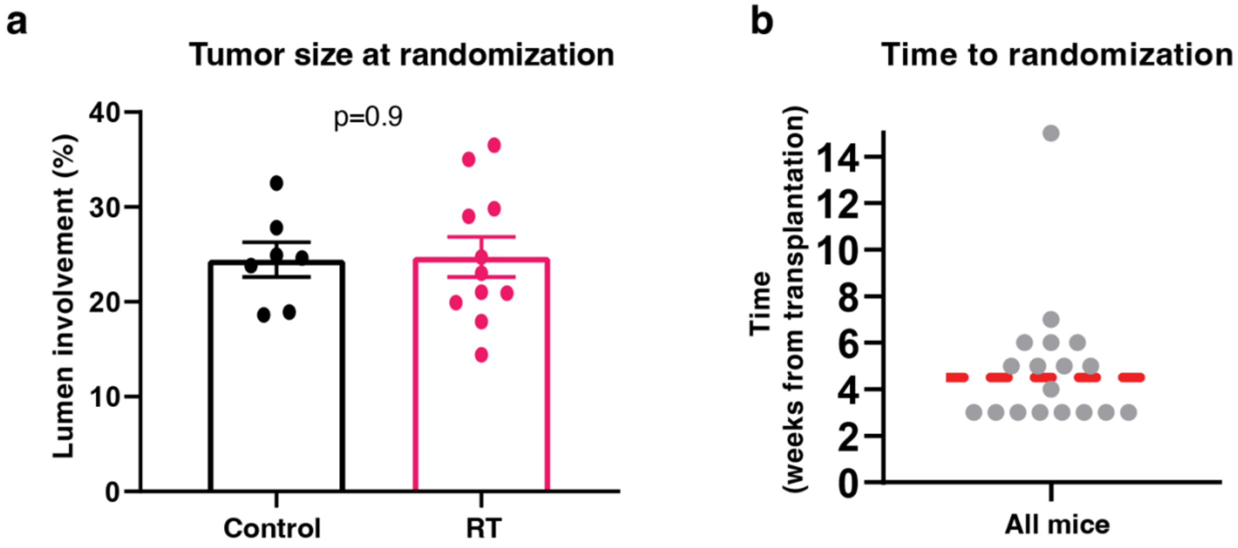


**Supplementary Figure 2. Tumor size and the time to randomization.** (**a**) Tumor size of the mice from the control (n=7) and RT groups (n=11) at randomization. Error bars denote the standard error of mean. Comparisons by unpaired two-tailed t-test. (**b**) Time from tumoroid transplantation until tumors involve 20-30% of the rectal lumen. The median time (4.5 weeks from transplantation) is indicated by a red line.

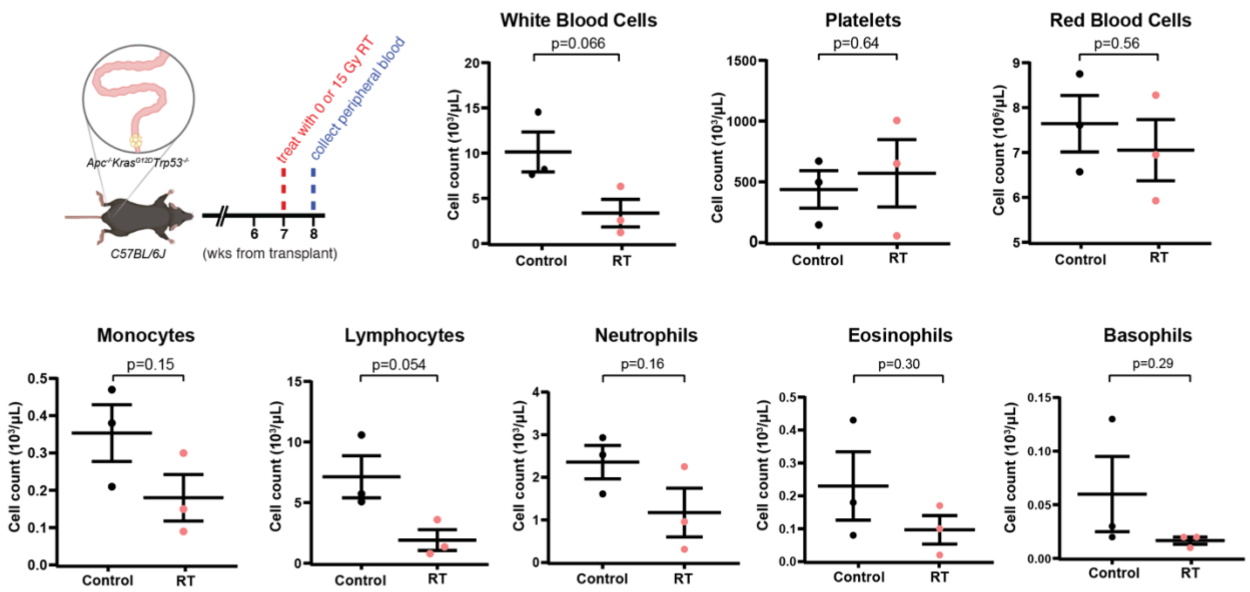


**Supplementary Figure 3. Complete blood count of peripheral blood from 3 control mice and 3 RT mice.** Control mice were not irradiated; RT received 15 Gy of pelvic radiation. Horizontal lines in the plots denote the mean and the standard error mean. P-values were provided by unpaired two-tailed t-test.

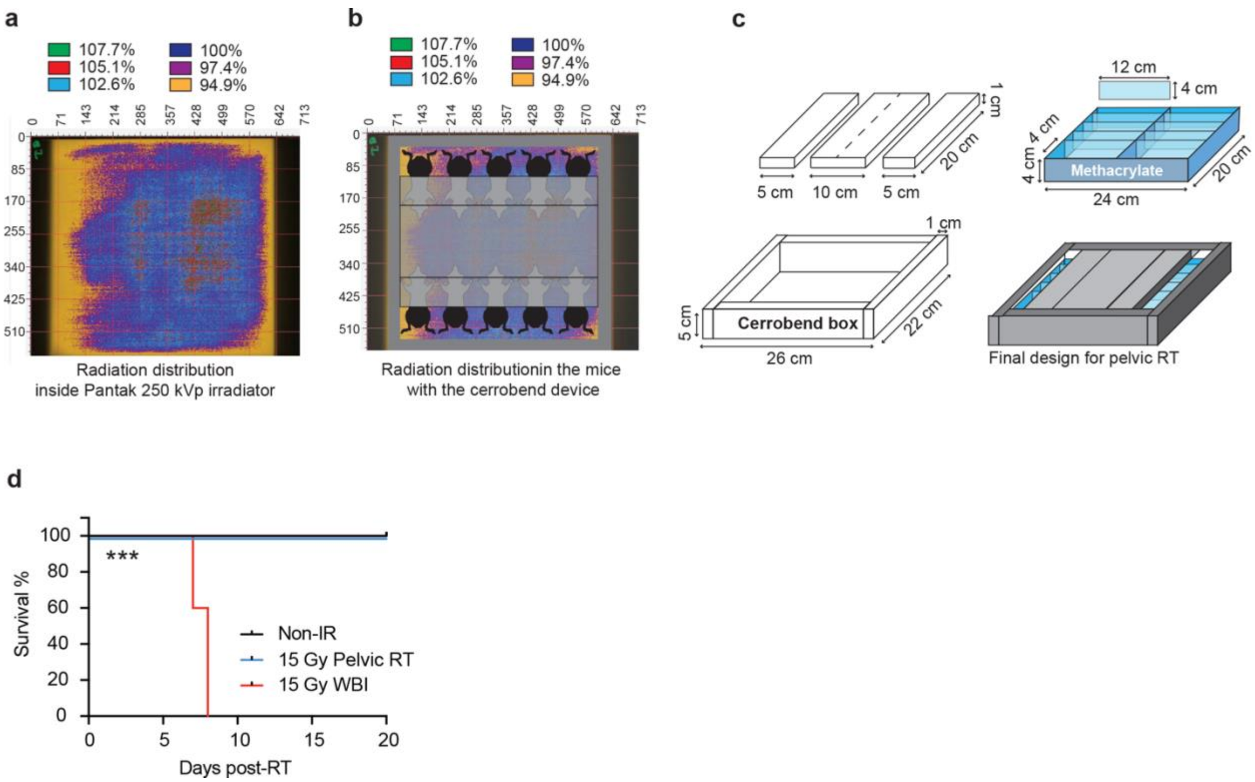


**Supplementary Figure 4. Customized cerrobend apparatus permits localized pelvic radiation.** (a) Relative dose by radiochromic film measurements[1] where 100% cooreponds to 15Gy. (b) Mouse position within customized cerrobend apparatus where 1cm thick cerrobend blocks can be positioned to shield the mouse head, thorax, and abomen leaving the pelvis exposed. Radiochromic film measurement confirmed shielded areas received less than 0.4% of prescribed dose. (c) Schematic and dimensions of customized cerobend apparatus with positioning for localized pelvic RT. (d) Mouse survival after 15Gy whole body irradiation as compared to 15Gy localized pelvic radiation. ***P<0.001.

**Supplementary Table 1. Colon and rectal cancer mouse models by anatomic relevance**

| Anatomic location | Tumorigenesis method | Advantages | Disadvantages | Engraftment rate | Ref |
|---|---|---|---|---|---|
| **Heterotopic Methods** | | | | | |
| Subcutaneous flank | Tumor injection | · Technically simple | · Ectopic microenvironment<br>· Immunocompromised host<br>· Does not metastasize | 25%, 81%[2]<br>56%[3] | [2,3] |
| Abdominal organ (Renal capsule, peritoneum) | Tumor injection | · High engraftment | · Ectopic microenvironment<br>· Immunocompromised host | | [4,5] |
| **Orthotopic Methods—Colon** | | | | | |
| Cecum | Surgical submucosal implantation of tumor | · High engraftment<br>· Immunocompetent host | · Not as anatomically relevant as a rectal cancer model<br>· Requires surgical technical expertise | 100%[6] | [6] |
| Predominantly colon | Colitis-induced | · Inexpensive<br>· Can be used in mice with any genetic background | · Mimics colitis-associated colorectal cancer, not sporadic colorectal cancer<br>· Tumor location is not specific to rectum<br>· Slow tumor growth (10-30 wks until tumor formation) | 38-100%[7]<br>80-100%[8] | [7,8] |
| Predominantly colon | Genetically modified murine model | · Mimics sporadic cancer<br>· Immunocompetent host | · Tumorigenesis in multiple organs; location not specific to rectum<br>· Expensive as it requires genetically engineered mice | 20-50%[9] | [9,10] |
| Distal colon | Genetically modified murine model with focal activation of mutations | · Mimics sporadic cancer<br>· Metastasis reported | · Expensive as it requires genetically engineered mice<br>· Slow tumor growth (≥10 weeks until >20% lumen obstruction) | 71-96%[11]<br>34-92%[12] | [11,12] |
| Distal colon | Colonoscopy-based mural injection of tumor | · High engraftment<br>· Metastasis reported | · Technically challenging<br>· Tumor does not initiate from the mucosa | 83%%[13]<br>90-92%[14] | [13,14] |
| **Orthotopic Methods—Rectum** | | | | | |
| Distal colon and rectum | Colitis induction + tumor enema | · Tumors grow from the mucosal lumen<br>· Metastasis reported<br>· Applicable to immunocompetent host | · Chemically induced colitis alters the tumor microenvironment<br>· Multifocal tumor formation along the colorectum<br>· Tumorigenesis: weeks to months | 100%[15]<br>20-100%[16]<br>94%[17]<br>62%[18] | [15–19] |
| Rectum | Intrarectal tumor injection | · Tumors form in distal rectum (1-2 mm above anus)<br>· High engraftment<br>· Applicable to immunocompetent host | · Tumor does not initiate from the mucosa<br>· Tumors are too close to the anus for endoluminal monitoring, so they need to be tagged with bioluminescent markers for imaging | 100%[20] | [20,21] |

#### Step-By-Step Procedures

##### Culturing and Passaging Mouse Rectal Tumor Organoids. (TIMING: 2-3 weeks)

Before starting cultivation, prepare enough EN organoid culture medium and thaw growth factor reduced Matrigel overnight at 4°C.

1. Culture tumor organoids (tumoroids) in four 50-μL Matrigel droplet domes per well in a 6-well plate. Add 2 mL of EN organoid culture medium per well. Exchange the medium every 2-3 days. Maintain subconfluent density.

**TIP:** Healthy tumoroids have an empty lumen lined with a thin bright layer without dark, necrotic areas.

2. When tumoroids approach confluence, carefully aspirate the medium from each well without disturbing Matrigel. We recommend using a P1000 pipette and not vacuum suction to avoid aspirating the Matrigel plugs.
3. If the tumoroids were cultured in a medium other than EN organoid culture medium, rinse each well twice with 2 mL of ice-cold PBS.
4. Add 2 mL of TrypLE Express Enzyme to each well, pipetting up and down with P1000 to break up Matrigel. Transfer two wells of dissociated solutions (approximately 4.5 mL) from the 6-well plate into a 15-mL conical tube.
5. Incubate the conical tube containing tumoroids in a 37°C water bath for 5-10 min.

**CRITICAL:** Incubate in the water bath long enough to dissolve Matrigel. Undissolved Matrigel will be spun down along with the tumoroids in step 7 and may affect tumoroid resuspension.

6. Add twice the volume of cold Basal medium (9 mL) to the tube and mechanically dissociate tumoroids by pipetting up and down 10-20 times in a 10 mL serological pipette.

**TIP:** Tumoroid dissociation rate is dependent on the incubation time with TrypLE Express Enzyme and the amount of pipetting.

7. Centrifuge the tube at 450g for 5 min at 4°C.
8. Aspirate the medium and resuspend the tumoroids in an appropriate volume of Matrigel.
9. Apply four 50 μL tumoroid-Matrigel droplets in each well and allow this to solidify in a 37°C incubator for 15 min.
10. Prepare an appropriate volume of EN organoid culture medium with 10 μM Y27632 and warm up to 37°C in a water bath.
11. Add 2 mL of EN organoid culture medium (+ Y27632) to each well.
12. Incubate the tumoroids in a 37°C incubator and replace the EN organoid culture medium (without Y27632) every 2-3 days.

##### Preparation of Mouse Rectal Tumoroids for Transplantation. (TIMING: 1-2 hours)

13. Carefully aspirate the medium from each well without disturbing Matrigel.

14. If the tumoroids were cultured in a medium other than EN organoid culture medium, rinse each well twice with 2 mL of ice-cold PBS.

15. Add 2 mL of TrypLE Express Enzyme to each well, pipetting with P1000 to break up Matrigel. Transfer two wells of dissociated solutions (approximately 4.5 ml) in a 6-well plate into a 15-mL conical tube. Dissociate 8 wells in total to make 4 tubes.

16. Incubate the conical tubes containing tumoroids in a 37°C water bath for 5-10 min.

**CRITICAL**: Incubate in the water bath long enough to dissolve Matrigel. Undissolved Matrigel will be spun down along with the tumoroids in step 19 and may affect tumoroid resuspension.

17. Add twice the volume of cold Basal medium (9 mL) to each conical tube and dissociate tumoroids by pipetting up and down 5-10 times in a 10-mL serological pipette.

18. Put the conical tube containing dissociated tumoroids on ice and repeat steps 13-17 for another 6 wells.

19. Centrifuge the tube at 450g for 5 min at 4°C.

20. Aspirate the medium and resuspend the tumoroids in 10 mL of Basal medium by carefully pipetting 10-20 times in a 10 mL serological pipette.

21. Place a 100-micron cell strainer on top of a 50-mL conical tube and filter the tumoroids to remove large aggregates.

22. Take 30 μL of tumoroids into a 0.2-mL tube and add an equal volume of trypan blue solution. Determine the number of viable tumoroids with a hemocytometer.

**TROUBLESHOOTING** (also shown in Supplementary Table 2): Counting the number of viable cells can be difficult if the tumoroids are not fully dissociated. Pellet down the tumoroids again at 450g for 5 min at 4°C. Resuspend the tumoroids in 500 μL of TrypLE Express Enzyme and incubate in a 37°C water bath for 5-10 min. Add three times the volume of cold Basal medium to a conical tube and dissociate tumoroids by pipetting 10-20 times using a P1000 pipette. Determine the number of viable tumoroids as outlined in step 22.

23. Calculate the total number of the tumoroids needed for transplantation.

24. Transfer the appropriate volume of tumoroids into a new 15-mL conical tube.

25. Centrifuge the tube at 450g for 5 min at 4°C.

26. Resuspend the tumoroids in an appropriate volume of ice-cold Tumoroid Transplantation medium to concentrate $2x10^5$ cells/50 μL per mouse. Place tumoroids on ice.

**Preparation of Mice (TIMING: 10 min)**

Refer to **Supplementary Video** 1 for a detailed visual of the steps for Preparation of Mice and Tumoroid Transplantation Procedure.

27. Anesthetize 6- to 8-week-old female C57Bl/6J mice using 2% isoflurane and oxygen in an induction chamber and then maintain proper anesthetic depth via nose cone.

**TROUBLESHOOTING:** If there is difficulty with achieving anesthesia, check if the isoflurane or oxygen supply is low. Replace any insufficient supplies.

**TROUBLESHOOTING:** If a mouse is over-anesthetized, reduce the inflow of isoflurane. Warm the mouse under a heating lamp if it is cold.

28. Place mice in sternal recumbency on a padded surface and flush out stool from the rectum using a thin, rounded gavage needle and syringe with approximately 5 mL of warm PBS (**Figure 1b. i**).

**Tumoroid Transplantation Procedure (TIMING: 3 min)**

29. Trim off a nonfiltered P200 pipette tip with a narrow opening so that the brush will fit through. This will serve as a guide for the brush to protect the anus from mechanical irritation.

**CRITICAL:** Make sure the brush will fit through the distal end of the pipette tip. Inserting the brush without the pipette tip as a guide will cause severe injury to the mouse anus, leading to bleeding and death. Use sufficient Vaseline to lubricate the anus.

30. Lubricate the guide with Vaseline and insert into the anus of the mouse (**Figure 1b. ii**).

**CRITICAL:** The pipette tip guide is only to protect damage to the anus. Do not insert the guide too far into the rectum as that will prevent injury to the distal rectum.

31. Insert the small-caliber brush through the anus and mechanically disrupt the rectal mucosa by gently pulling the brush out and in 2-5 times (**Figure 1b. iii**). Use your wrist to gently twist the brush as you pull it out and in. We inserted only about 5-10 mm of the brush beyond the tip of the guide, as strong resistance beyond this depth will be encountered and lead to significant rectal injury. Please note that excess force and irritation can lead to perforation of the colon and death. Wipe the tip of the brush on a white napkin and look for a red/pink tinge to confirm blood on the brush tip (**Figure 1b. iv**).

**CRITICAL**: Do not force the brush into the rectum if you encounter resistance, as this can lead to bowel perforation.

**TROUBLESHOOTING:** If there is resistance when trying to insert the brush, this may be due inadequate trimming of the tip in step 29 or an acute angle of the P200 pipette tip guide. Take out the P200 pipette tip guide and ensure that the distal tip is large enough for the brush to fit through. Trim more of the tip if needed. Adjust the angle of the P200 pipette tip guide so it is horizontal. Better aligning the guide with the rectal lumen will allow the brush to easily glide in.

32. After adequate mechanical injury to the mucosa has been achieved, remove the P200 pipette tip.

33. Lift the mouse at the base of the tail so that the rectal cavity is vertical (**Figure 1b, v**).

34. Instill 50 μL of tumoroids prepared in Tumoroid Transplantation medium into the rectum using a P200 pipette (**Figure 1b, vi**).

**TROUBLESHOOTING**: Depositing the tumoroids too quickly may induce colorectal peristaltic activity and result in spillage of the contents. Pipetting the tumoroids slowly will reduce this occurrence. If spillage occurs, pipette up the remaining contents and re-instill them slowly. This may require a few repetitions to instill the tumoroids when the colorectum is not in active peristalsis.

35. Immediately coat the anus with 5-10 µL of Vetbond adhesive and pinch the anus closed to prevent luminal contents from being excreted (**Figure 1b, vii-viii**).
36. Keep lifting the tail of the mouse to allow the Vetbond to dry for 1 min to prevent spillage.
37. Allow the mouse to recover under a heat lamp.
38. After 6 hours, remove the Vetbond by gently wiping the anus with a wet gauze pad.

**Tumor monitoring by colonoscopy/endoscopy (TIMING: 10 min)**

Tumors are usually detected starting at 2-4 weeks after transplantation. We recommend surveillance with colonoscopy every 1-2 weeks after the second week after transplantation.

**CAUTION**: Early colonoscopy (<2 weeks after transplantation) can result in intestinal perforation, because the mouse rectum is still friable and healing from the injury

39. Anesthetize mice using 2% isoflurane and oxygen in an induction chamber and then maintain proper anesthetic depth via nose cone.

**TROUBLESHOOTING**: If there is difficulty with achieving anesthesia, check if the isoflurane or oxygen supply is low. Replace any insufficient supplies.

**TROUBLESHOOTING**: If the mouse is over-anesthetized, reduce the inflow of isoflurane. Warm the mouse under heating lamp if it is cold.

40. Place mice in sternal recumbency on a padded surface and flush out stool from the rectum using a thin, rounded gavage needle and syringe with approximately 5 mL of warm PBS.

**TROUBLESHOOTING**: If no stool is removed, flush out the rectum with more PBS. Massage the abdomen until stool is expelled.

41. Insert a 1.9-mm rigid bore endoscope and insufflate. Using a 300W Xenon light source, capture a video of the procedure.

**CRITICAL**: To prevent colonic perforation, insufflation must be used sparingly.

42. Recover mice in the cages under a heated lamp.
43. Export colonoscopy videos and take snapshots of the tumors for measurement as described[44].

**Rectal Irradiation (TIMING: 15 min)**

44. Weigh mice and give appropriate ketamine/xylazine anesthetic (100 mg/mL; 10 µl/g body weight) via intraperitoneal injection.

**CRITICAL:** After ketamine/xylazine administration and completion of irradiation treatment, mice require up to one hour of monitoring for proper recovery. As ketamine is a controlled substance, this anesthetic cocktail must be maintained in a stationary, double-locked cabinet as required by United States Drug Enforcement Agency regulations. Detailed records of use must also be maintained as required by institutional policy.

45. After proper anesthetic depth has been achieved, place mice in dorsal recumbency into the customized lead apparatus, keeping only the distal abdominal region exposed (**Figure 2a**). Limit offsite radiation exposure with additional protective lead strips (**Figure 2b**).

46. Set the X-Ray tube maximum potential to 225 kV/12.5 mA. Administer a single 15-Gy dose of ionizing radiation at a rate of 117.5 cGy/min.
47. Remove mice from the X-ray system and place into their cages under a heated lamp for recovery.

**CRITICAL**: Weigh mice at the beginning of the experiment and weigh them every other day for the first week after irradiation and thereafter every other week to identify signs of sickness.

#### REAGENTS

##### Tumoroid culture

- Matrigel, Growth factor reduced (Corning, cat. no. 356231): used for tumoroid culture
- Matrigel, Basement (Corning, cat. no. 356237): used for endoluminal transplantation
- Advanced DMEM/F12 (Thermo Fisher Scientific, cat. no. 12634-010)
- Penicillin–streptomycin, 10,000 U/mL (Thermo Fisher Scientific, cat. no. 15140-122)
- HEPES, 1 M (Quality Biological, cat. no. 118-089-721)
- GlutaMAX supplement, 100× (Thermo Fisher Scientific, cat. no. 35050-061)
- N-acetyl-L-cysteine (Sigma-Aldrich, cat. no. A9165)
- Nicotinamide (Sigma-Aldrich, cat. no. N0636)
- Bovine serum albumin (BSA) (Sigma-Aldrich, cat. no. A2058)
- Recombinant mouse EGF (mEGF; Thermo Fisher Scientific, cat. no. PMG8043)
- Dulbecco's Modified Eagle's Medium (DMEM; ATCC, cat. no. 30-2002)
- Fetal Bovine Serum (FBS; Sigma, F2442)
- B27 Supplement (Thermo Fisher Scientific, cat. no. 17504-044)
- Y27632 dihydrochloride (ROCK inhibitor; Sigma-Aldrich, cat. no. Y0503)
- TrypLE Express Enzyme (Thermo Fisher Scientific, cat. no. 12605-010)
- Cell Recovery Solution (Corning, cat. no. 354253)
- PBS (Corning, cat.no. 21-031-CV)
- 0.4% Trypan Blue solution (Sigma, cat.no. T8154)

##### Immunofluorescence

- Rat anti-mouse F4/80-Alexa Fluor 594 monoclonal antibody, clone BM8 (Biolegend, cat.no. 123140) (1:100)
- Rabbit anti-mouse CD206 polyclonal antibody (Abcam, cat.no. ab64693) (1:1000)
- Armenian hamster anti-mouse CD11c monoclonal antibody, clone N418 (eBioscience, cat. no 14-0114-82) (1:100)
- Rabbit anti-mouse CD4 monoclonal antibody (Abcam, cat.no. ab183685) (1:200)
- Rat anti-mouse CD8 monoclonal antibody, clone 4SM15 (eBioscience, cat.no. 14-0808-80) (1:100)
- Rat anti-mouse FOXP3 monoclonal antibody, clone FJK-16s (Invitrogen, cat.no. 14-5773-82) (1:100)
- Anti-Rat 594 fluorophore (Invitrogen, cat.no. A-21209) (1:500)
- Donkey anti-rabbit secondary antibody, Alexa Fluor 488 (Invitrogen, A-21206) (1:500)
- Donkey anti-rabbit secondary antibody, Alexa Fluor 647 (Invitrogen, A-31573) (1:500)

- Goat anti-Armenian hamster secondary antibody, Alexa Fluor 488 (Invitrogen, A-21110) (1:500)
- Rat IgG2a-Alexa Fluor 594 isotype control (Biolegend, cat.no. 400555) (1:100)
- Rabbit IgG isotype control (Abcam, cat.no. ab172730)
- Rat IgG2a isotype control (R&D, cat.no. MAB006)
- Armenian Hamster IgG isotype control (eBioscience, cat.no. 14-4888-81)
- Hoescht 33342 (Sigma, B2261, dissolved in distilled water to 1 mg/mL) (1:1000)

**EQUIPMENT**

**Common equipment**

- Humidified $CO_2$ incubator (37 °C, 5% $CO_2$) (NuAire, model no. NU-8700)
- Biosafety cabinet, class II type A2 (NuAire, model no. S407-400)
- Light microscope for tissue culture (Nikon, TMS model no. 213021)
- Water bath (37°C) (Boekel Grant, model no. PB-1400)
- Benchtop centrifuge (Eppendorf, model no. 5810R)
- Cell culture dish, 150 mm (Falcon, cat.no. 353025)
- Suspension cell culture plate, 6-well (Greiner Bio-One, cat. no. 657185)
- Conical tube, 15 mL (VWR, cat. no. 525-1069)
- Conical tube, 50 mL (VWR, cat. no. 525-1074)
- 0.2-mL tube (Fisherbrand, cat. no. 14-230-225)
- Microcentrifuge tube, 1.5 mL (Crystalgen, cat. no. L2052): Autoclave and dry
- 5-mL tube (Axygen, cat. no. MCT-500-C): Autoclave and dry
- Pipette aid (BrandTech, model no. accu-jet pro 26333)
- Serological pipettes (5, 10 and 25 ml) (Falcon, cat. no. 357543, 357551, 357535)
- Suction pipette (Falcon, cat. no. 357558)
- Micropipettes (P20, P200 and P1000) and pipette tips (Gilson, Crystalgen)
- 500-mL bottle-top vacuum filter, 0.22 µm (Corning, cat. no. 431118)
- Hemocytometer (Electron Microscopy Sciences, cat. no. 63511-13)
- 60-mL syringe (BD, cat. no. 309653)
- 0.45-µm syringe filter (Thermo Scientific, cat. no. 723-2545)
- 100-µm Cell Strainer (BD Falcon, cat. no. 352360)
- Glass microscope slide (Fisher Scientific, cat. no. 12-544-7)
- Glass microscope cover slide (Fisher Scientific, cat. no.12542B)

**Mechanical brush injury to the rectal mucosa**

- Dissociated tumoroids in 5% Matrigel-PBS
- Small caliber nylon brush (2.5 mm) (Karl Storz #27650C)
- Nonfiltered P200 pipette & tip
- Oral gavage needle (Roboz, cat. no. FN 7905)
- PBS (Corning, cat. no. 21-031-CV)
- 10-mL syringe (BD, cat. no. 309604)
- Isoflurane (available through animal facility)
- Vaseline
- Vetbond Tissue Adhesive (3M, cat. no. 1469SB)

**Colonoscopy/Endoscopy**

- 1.9-mm rigid 30° small animal endoscope (Karl Storz, part 64301BA)
- Oral gavage needle (Roboz, cat. no. FN 7905)
- 10-ml syringe (BD, cat. no. 309604)
- PBS (Corning, cat. no. 21-031-CV)
- Image J software (NIH)

**Rectal Irradiation**

- X-Rad 320 machine (Precision X-Ray)
- Customized cerrobend block (provided by Memorial Sloan Kettering Cancer Center Machine Shop Facility; fabrication method available upon request)
- Ketamine/Xylazine anesthetic: 2 mL of 100 mg/mL Ketamine, 1 mL of 20 mg/mL Xylazine, 17 mL $H_2O$ (available through animal facility)

**Complete Blood Count of Peripheral Blood**

- Hemavet 950 (Drew Scientific Serial No. 03641)
- BD Microtainer Tubes with K2E (K2EDTA) (Becton, Dickinson, and Company; Reference Number 365974).
- Stuart Roller Mixer SRT9D

#### REAGENT SETUP

Bovine serum albumin (BSA)
Dissolve 5 mg of BSA in 50 mL of PBS to make 10% (wt/vol) stock solution. Filter-sterilize the solution with a 0.45-µm syringe filter into a sterile 50-ml conical tube. Aliquot and store at -20 °C.

Mouse epidermal growth factor (mEGF)
Dissolve 1 mg of mEGF in 2 mL of PBS with 0.1% (wt/vol) BSA to make a 500 µg/mL stock solution. Aliquot and store at -20°C.

N-acetyl-L-cysteine (NAC)
Dissolve 816 mg of N-acetyl-L-cysteine in 10 mL of water to make a 0.5 M stock solution and filter-sterilize the solution with a 0.45-µm syringe filter into a sterile 50-mL conical tube. Aliquot and store at -20 °C.

Nicotinamide
Dissolve 6.11 g of nicotinamide in 50 mL of water to make a 1 M stock solution and filter-sterilize the solution with a 0.45-µm syringe filter into a sterile 50-mL conical tube. Aliquot and store at -20 °C.

ROCK inhibitor; Y-27632
Dissolve 5 mg of ROCK inhibitor (Y-27632) in 1.56 mL of PBS to make a 10 mM stock solution. Aliquot small volumes (50 µL) and store at -20 °C.

Collection medium
Add 5 mL of penicillin–streptomycin (100X), 5 mL of HEPES (1 M), 5 mL of GlutaMAX (100X), 1 mL of NAC (0.5 M), and 10 mL of B27 supplement (50X) into 474 mL Advanced DMEM/F12 to make up 500 mL of medium. Freshly prepare the medium before use.

Noggin conditioned medium
Maintain 293T-Noggin cells in DMEM supplemented with 10% FBS. To make Noggin conditioned medium, split cells into 150-mm culture dishes and grow until confluent. Rinse each dish 1-2 times with warm PBS and add 30 mL of Collection medium. After 1 week, collect this conditioned medium into 50-mL conical tubes. Centrifuge the tubes at 800 G for 5 min at 4°C and filter the supernatant through a 0.22 µm bottle-top vacuum filter. Aliquot 5 mL into 15-mL conical tubes and store at -20 °C.

Basal medium
Add 5 ml of penicillin–streptomycin (100X), 5 ml of HEPES (1 M), 5 ml of GlutaMAX (100X), and 1 ml of NAC (0.5 M) into 484 ml Advanced DMEM/F12 to make up 500 ml of medium. Store at 4°C for up to 1 month.

EN organoid culture medium
Mix 44 mL of Basal medium, 500 µL of BSA (10%, wt/vol), 500 µL of nicotinamide (1 M), 5 mL of Noggin conditioned medium, and 5 µL of mEGF (500 µg/mL, wt/vol) to make up 50 mL of medium. Store at 4°C for up to 2 weeks.

Tumoroid Transplantation medium
Dilute 50 µL of Matrigel (Basement) with 950 µL of cold PBS to make 5% (vol/vol) Matrigel solution.

**Supplementary Table 2. Troubleshooting Table**

| Step | Problem | Possible reasons | Solution |
|---|---|---|---|
| 22 | Viable tumoroid count is inconsistent with the relative cell confluency | Counting the number of viable cells can be difficult if the tumoroids are not fully dissociated | Pellet down the tumoroids again at 450g for 5 min at 4°C. Resuspend the tumoroids in 500 µL of TrypLE Express Enzyme and incubate in a 37°C water bath for 5-10 min. Add three times the volume of cold Basal medium to a conical tube and dissociate tumoroids by pipetting 10-20 times using a P1000 pipette. Determine the number of viable tumoroids as outlined in step 22. |
| 27, 39 | Difficulty with achieving anesthesia | Low supply of isoflurane or oxygen | Replace any insufficient supplies. |
| 27, 39 | Mouse is over-anesthetized | Isoflurane setting is too high | Reduce the inflow of isoflurane. If the mouse is cold, warm it under a heating lamp. |
| 31 | Resistance when trying to insert the brush | The P200 pipette tip may be too small to allow entry of the brush<br><br>The angle of the P200 pipette tip guide may be too acute | Take out the P200 pipette tip guide and ensure that the distal tip is large enough for the brush to fit through. Trim more of the tip if needed. Adjust the angle of the P200 pipette tip guide so it is horizontal. Better aligning the guide with the rectal lumen will allow the brush to easily glide in. |
| 34 | Immediate spillage of the instilled tumoroids | Depositing the tumoroids too quickly may induce colorectal contraction | Pipetting the tumoroids slowly will reduce this occurrence. If spillage occurs, pipette up the remaining contents and re-instill them slowly. This may require a few repetitions to instill the tumoroids when the colorectum is not contracting. |
| 40 | No stool removed | Inadequate flushing | Flush out the rectum with more PBS. Massage the abdomen until stool is expelled. |
